# Flanker task-elicited Event Related Potential sources reflect human recombinant Erythropoietin differential effects on Parkinson’s patients

**DOI:** 10.1101/864397

**Authors:** Maria L Bringas Vega, Shengnan Liu, Min Zhang, Ivonne Pedroso Ibañez, Lilia M. Morales Chacon, Lidice Galan Garcia, Vanessa Perez Bocourt, Marjan Jahanshahi, Pedro A Valdes-Sosa

## Abstract

We used EEG source analysis to identify which cortical areas were involved in the automatic and controlled processes of inhibitory control on a flanker task and compared the potential efficacy of recombinant-human erythropoietin (rHuEPO) on the performance of Parkinson’ s Disease patients.

The samples were 18 medicated PD patients (nine of them received rHuEPO in addition to their usual anti-PD medication through random allocation and the other nine patients were on their regular anti-PD medication only) and 9 age and education-matched healthy controls (HCs) who completed the flanker task with simultaneous EEG recordings. N1 and N2 event-related potential (ERP) components were identified and a low-resolution tomography (LORETA) inverse solution was employed to localize the neural generators.

Reaction times and errors were increased for the incongruent flankers for PD patients compared to controls. EEG source analysis identified an effect of rHuEPO on the lingual gyri for the early N1 component. N2-related sources in middle cingulate and precuneus were associated with the inhibition of automatic responses evoked by incongruent stimuli differentiated PD and HCs.

From our results rHuEPO, seems to mediate an effect on N1 sources in lingual gyri but not on behavioural performance. N2-related sources in middle cingulate and precuneus evoked by incongruent stimuli differentiated PD and HCs.

## Introduction

Discovering neuroprotective agents to slow down the progression of Parkinson’s Disease (PD) and, importantly, to improve cognitive deficits is an active area of research (Athauda & Foltynie, 2015). The search for agents to supplement usual dopaminergic treatments directed towards motor symptoms is not surprising since the characteristic motor impairment of patients is usually accompanied by cognitive deficits (Kehagia, Barker, & Robbins, 2010). Since cognitive dysfunction has a negative impact on the quality of life of patients(Schrag, Jahanshahi, & Quinn, 2000); finding effective therapies that target cognition in PD is of paramount importance. As an example, we found that human recombinant erythropoietin (rHuEPO) (Pedroso et al., 2012) improved general measures of cognition in chronically medicated PD patients, an additional benefit to that obtained on their usual medical treatment. This result extends to PD the evidence for neuroprotective properties of rHuEPO already described in other neurologic diseases (Brines & Cerami, 2005) and is supported by the anti-apoptotic, anti-inflammatory and cytoprotective effects of EPO in PD animal models (Sirén, Faßhauer, Bartels, & Ehrenreich, 2009)(Xue, Zhao, & Guo, 2007). This promising result suggested the need to further study the effect of rHuEPO on cognition in PD.

We believe that to further understand the effect of rHuEPO on cognition in PD patients we need to examine its effect on specific stages of information processing. This is because the overt behavioural measures used in our previous study: a) do not have temporal sensitivity, being the end outcome of many sequential processes, and b) do not reflect localized neural activity. Consequently, and as a first objective, we zeroed in on very early automatic neural processes involved in inhibitory control, the lack of which is so common in non-demented PD patients. This early lack of inhibitory control is easily measured in a number of tasks such as the Stop signal, go no-go, Stroop, Hayling Sentence Completion task and the Simon task described in (Obeso et al., 2011) and (Seer, Lange, Georgiev, Jahanshahi, & Kopp, 2016). However, we decided to use a very well-studied paradigm: Ericksen’s Flanker Task (Eriksen & Eriksen, 1974). It explores the lack of inhibition related to the difficulty in suppressing interference by incongruent stimuli. It allows the evaluation of very short latency automatic activation to incongruent flankers around 100 msec. and other, controlled processes, around 200 msec. These produce increased reaction times (RTs) and errors in incongruent trials versus congruent trials in PD patients in comparison with normal (eg. (P Praamstra, Stegeman, Cools, & Horstink, 1998; Peter Praamstra, Plat, Meyer, & Horstink, 1999; S A Wylie et al., 2009; Scott A Wylie, Stout, & Bashore, 2005). It is, however, the early ERP responses that are of interest here, not the overt behavioural response indexed by the RT which occurs later about 400 msec.

There is no clear way to study these early responses behaviourally. However, these processes might be probed by direct measurements of fast neural responses such as those provided by event-related responses (ERPs). In particular, the Flanker task elicits the N1, N2 and P3 ERP components, which are related to automatic and controlled process respectively (Pires, Leitai, Guerrini, & Simoes, 2014). Here, we will focus only on the early components N1 and N2. The N1 component has not been, to our knowledge sufficiently studied in the Flanker task in PD. However, the fronto-central N2 on incongruent trials of flanker tasks in patients with PD have received more attention (M Falkenstein, Willemssen, Hohnsbein, & Hielscher, 2006; J. R. Folstein & Van Petten, 2008; Verleger et al., 2010; S A Wylie et al., 2009; Scott A Wylie et al., 2005). The comparison of medicated PD patients and drug-naïve de novo PD patients showed that neither the presence of PD (see also (Verleger et al., 2010) nor dopaminergic medication modulates N2 amplitude variability on incongruent conditions of flanker tasks (for a discussion see a review of ERP and cognition in PD by Seer et al., 2017). It seems logical then to determine if the additional cognitive improvement produced by rHuEPO with respect to dopaminergic treatment, is accompanied by changes in the early components in the N1 and N2 ERP components, helping us to pinpoint one of the stages of cognitive processing affected by this drug. Furthermore, in addition to finer grained timing information, it is possible to leverage source localization methods to identify the neural sources of any ERP component change.

Therefore, the aim of our study is to use a flanker task to identify if rHuEPO improves automatic and controlled inhibitory control in PD patients and to locate the neural generators of these processes. This could be a first step in identifying an ERP biomarker for this type of cognitive process to be used in clinical trials.

## Materials and Methods

## Methods

### Description of the Sample and Clinical Trial

Eighteen PD patients (Hoehn and Yahr stages I to III, mean age 53.9, SD 3.2 years) were recruited at the Clinic of Movement Disorders and Neurodegeneration, Centro International de Restauracion Neurologica (CIREN) in La Habana, Cuba to participate in a safety clinical assay of Erythropoietin (rHuEPO) in PD. The design of this investigation, results, scheme of application and doses employed may be found in (Pedroso et al., 2012). Inclusion criteria were a clinical diagnosis of idiopathic PD according to the UK Brain Bank criteria and a good response to dopaminergic treatment and aged between 45-75 years (Hughes, Daniel, Kilford, & Lees, 1992). Exclusion criteria were manifestation or indicative signs of major cognitive impairment, psychotic symptoms, and/or presence of other chronic diseases. Nine of the PD patients, through random allocation, received additionally to their usual anti-parkinsonism medication, rHuEPO for five weeks and the other nine did not. rHuEPO approved and registered for use in humans was obtained at the Centro de Inmunologia Molecular, La Habana Cuba (ior® EPOCIM). There were no significant differences in age, years of education or duration of illness between the two PD groups. To exclude dementia and major depression, the Mini Mental State Examination and the Hamilton Depression Scale were respectively administered (M. F. Folstein, Folstein, & McHugh, 1975; Hamilton, 1960). All patients were assessed on the motor subscale of the Unified Parkinson’s Disease Rating Scale (UPDRS) both during “on” (mean 6.3, SD1.1) and “off” medication (mean 21.7, SD 4.3) states.

For the purpose of comparisons, 9 healthy controls (HCs) matched in age (mean 51.2, SD 3.9 years) and educational level were recruited at the same clinic. The PD patients were tested on their usual anti-parkinsonism medication. The patients and controls signed an informed consent to participate in this study as a complement of the clinical trial following the CIREN ethics committee regulations.

### Eriksen’s Flanker Task

All participants completed the Eriksen’s Flanker task, while the EEG was simultaneously recorded. Each trial of the task consisted of the presentation of a set of 5 ordered letters (HHHHH or SSSSS) for the congruent condition and 5 letters with H or S at the centre and different laterals or flankers (SSHSS or HHSHH) for the incongruent condition. Participants were instructed to respond to the central letter, whether H or S, by pressing a key with the index finger of the right or left hand respectively. Participants were instructed to respond as fast and as accurately as possible. A total of 480 trials in two blocks, each lasting 8 minutes were completed. In each block 80 stimuli were shown for the congruent condition and 160 for the incongruent. Only the correct responses with reaction times (RTs) >150 and <800 msec. were selected for analysis.

The physical characteristics of the stimuli were black letters on a white frame with a height = 1.5 cm. and Lenght= 7 cm., under 6 ^0^ a visual angle. The distance of the participant to the computer monitor was 60 cm. Each stimulus was presented at the centre of the screen for 190 msec., followed by a fixed interstimulus interval (ITI) of 1735 msec. A training block of 40 stimuli was designed to ensure task instructions were understood.

### ERP measurement

The Electroencephalogram (EEG) was continuously recorded at a sampling rate of 512 Hz from 64 electrodes located at standard positions of the International 10/20 System using a Brain Vision system (*https://www.brainproducts.com/products_by_apps.php?aid=5*) (Jasper, 1958). Linked ears were used as on-line reference and the front as earth. To monitor eye movement artefacts, the electro-oculogram (EOG, horizontal and vertical) was recorded from electrodes placed 1 cm to the left and right of the external canthi, and from an electrode beneath the right eye.

Data were filtered using 1-30 Hz and a notch filter to eliminate the 60Hz powerline artefact. All data were referenced using an average reference to all the channels. The baseline was corrected between −200 to – 0 msec. Epochs with electric activity exceeding baseline activity by 100 µV were considered as artefacts and were automatically rejected from further processing (15% of epochs related to hits and 11% of the epochs related to errors). For the analysis, several electrodes were excluded (EOG, ECG, TP9 and TP10).

ERPs were obtained from the EEG recordings for each participant for all the electrodes within the two experimental conditions and averaged over the two groups using Analyzer software (*https://www.brainproducts.com/productdetails.php?id=17).* Epochs of 800 msec. (from −200 msec. (baseline) until 600 msec. post-stimulus onset) were analyzed locked to the stimulus. We selected two windows to examine the stimulus-locked ERPs, using only the correct response averages for the N1 (80-180 msec.) and N2 (200-300 msec.) components in the expected time-windows (see ERPs guidelines in (Picton et al., 2000). Henceforth we will refer to these averages simply as the amplitude of the N1 and N2 components. The average waveform for each participant and each condition was estimated in all the electrodes, but the averaged waveform for group are plotted below for the electrode with the higher statistics amplitudes.

In order to localize the generators of the ERP components, a lead field was constructed for each participant to calculate the (volume-constrained) inverse solution, at the two selected latencies using LORETA (Low Resolution Tomography) (*http://www.uzh.ch/keyinst/loreta)*. (Pascual-Marqui et al., 1999)For LORETA, the intracerebral volume is partitioned into 6239 voxels at 5mm spatial resolution.

### Statistical analysis

We now summarize the experimental design. Our sample is divided into 3 **Groups**: 9 Parkinson patients with the usual treatment (PD Control), 9 patients with the usual treatment plus EPO (PD rHuEPO), and 9 healthy controls (HCs). Additionally, the ERPs for each participant was recorded in two conditions: congruent and incongruent.

For each participant the following variables were used in this paper:

1. Reaction Time and errors to the Flanker task
2. Amplitude of the N1 and N2 ERP component at the 60 EEG scalp electrodes.
3. Power of the N1 and N2 sources component for the 6239 source voxels.

The statistical analyses performed were:

a. Reaction Times and errors were analysed using a two-way repeated measure ANOVA with the with Group (HCs, PD Control and PD rHuEPO) as the between group factor and the experimental condition (incongruent versus congruent) as the within-subject repeated measures factor. We report the F statistic and the p value for tests of the main effect and the interaction. The Greenhouse-Geisser adjustment was applied since lack of sphericity was observed. These analyses were completed with STATISTICA 7.0.
b. An exploratory analysis of the differences in ERP amplitude topographies between the HCs and PD Control +PD rHuEPO groups was carried out by means of a multivariate t test that corrects for multiple comparisons by means of a permutation technique. The permutation test has the following advantages: the tests are distribution free that control the experiment-wise error for the simultaneous univariate comparisons, no assumptions of an underlying correlation structure are required, and they provide exact p-values valid for any number of subjects, timepoints and all 60 electrodes. The overall significance level was selected to be 0.05. The method is described in (Galán, Biscay, Rodríguez, Pérez-Abalo, & Rodriguez, 1997; Galan, Biscay, Valdes, Neira, & Virues, 1994) as implemented in the software NEEST from Neuronic *http://www.neuronicsa.com/*. This allowed the selection of a:
  1. A subset of electrodes to be subjected to Multivariate Analysis of Variance (MANOVA) to be described in c) below.
  2. The selection of most representative electrodes to plot the N1 and N2 grand average ERPs.
  3. The analysis of time intervals to be further studied.
c. Examine for each ERP component, and for their selected group of electrodes, a repeated measures Multivariate Analysis of Variance (r-MANOVA) for the design Group by Condition with a significance level set at the 0.05 level. The different contrasts for the interaction and main effects were tested by using the Wilk’s lambda, approximated by an F function and the p value reported. Note that this allows a simultaneous confidence interval for contrasts on group differences and to examine which electrode contribute to the effects. The MANOVA was that implemented in the STATISTICA 7.0. package.
d. Further analysis for selected differences of the ERP component source images between selected groups was carried out using the LORETA-built-in voxel-wise randomization tests with 2000 permutations (Nichols & Holmes, 2001), based on statistical nonparametric mapping. Voxels with significant differences (p<0.01, corrected for multiple comparisons) between contrasted conditions were located with the coordinates of the AAL (Automated Anatomical Labelling of Activations) 116 structures atlas of the Montreal Neurological Institute (MNI) (Tzourio-Mazoyer et al., 2002).

## Results

### Behavioural results

#### Reaction time

The differences between the three groups were significant for Factor Group: (F(2,24)=7.47, p=0.003), the Condition was not significant as we predicted in the preliminary analysis (F(2,24)=3.22, p=0.06). The interaction of Group*Condition also was not significant (p>0.8). The contrast between the two groups of patients (PD Control and PD rHuEPO) didn’t show differences in the reaction time (F(2,15)=0.62, p=0.55). Table 1 shows the performance of the PD groups separately and Table 2 the fusion of PD patients versus HCs.

**Table 1:** The results of the reaction times and the percent errors for the congruent and incongruent trials for the PD patients with and without rHuEPO. The values in the table are means with standard deviations in parenthesis.

|  | PD rHuEPO n=9 |  | PD Control n=9 |  |
| --- | --- | --- | --- | --- |
|  | congruent | incongruent | congruent | incongruent |
|  | Means (SD) | Means (SD) | Means (SD) | Means (SD) |
| Reaction times msec. | 459.33 (71.76) | 479.89 (49.43) | 460.22 (72.10) | 488.22 (63.76) |
| Percent errors | 13.22 (7.76) | 43.22 (21.37) | 8.78 (6.76) | 32.00 (15.79) |

**Table 2:** The results of the reaction times and percent errors for the congruent and incongruent trials for the Parkinson’s. disease (PD) patients and healthy control (HCs) groups. The values in the table are means with standard deviations in parenthesis.

|  | (PD rHuEPO + PD Control) n=18 |  | HCs n=9 |  |
| --- | --- | --- | --- | --- |
|  | congruent | incongruent | congruent | incongruent |
|  | Means (SD) | Means (SD) | Means (SD) | Means (SD) |
| Reaction time msec. | 459.78 (69.79) | 484.06 (55.51) | 411.22 (52.00) | 431.33 (43.47) |
| Percent errors | 9.00 (3.81) | 37.61 (19.12) | 3.33 (2.40) | 11.00 (7.42) |

#### Errors

The differences between the errors in the three groups were significant for Factor Group: (F(2,15)=10.49, p=0.0014), and for Condition, (F(2,24)=11.6, p=0.0003), but not for the interaction Group*Condition (p=0.1). The comparison between the two PD groups were significant only for Condition, incongruent (F(1,16)=55.3, p=0.00001, and not for the congruent condition (F(1,16)=1.88, p=0.18).

When using the contrast comparing all PD patients and HCs (Table 2), the results were consistent with previous findings where the RTs increased with incongruent flankers compared to congruent for both groups.

### Exploratory results of ERPs

As mentioned in the Methods, the multivariate t tests corrected for multiple comparisons with permutation tests provides exact p-values, valid for any number of participants, timepoints and recording sites yielded as significant the ERP components in the midline at the 0.05 level. Within this group the most significant ERP was Oz for N1 and Cz for N2 as described in the literature. We will therefore concentrate on these electrode sets henceforth since they all are significant above the globally valid significance threshold.

The same procedure allows, additionally, to select the time windows and which factor (Condition or Group) to be further analyzed. Figure 1 illustrates, for one derivation, the statistics shown above the red line, the latencies with significance for each factor (Group or Condition) in all the time window for analysis. The interaction between them was not significant at any time. The exploratory analysis between experimental conditions did not reflect significant differences in the time range for the early ERP components N1 and N2 (around 100 and 200 msec. respectively).

**Figure 1.**
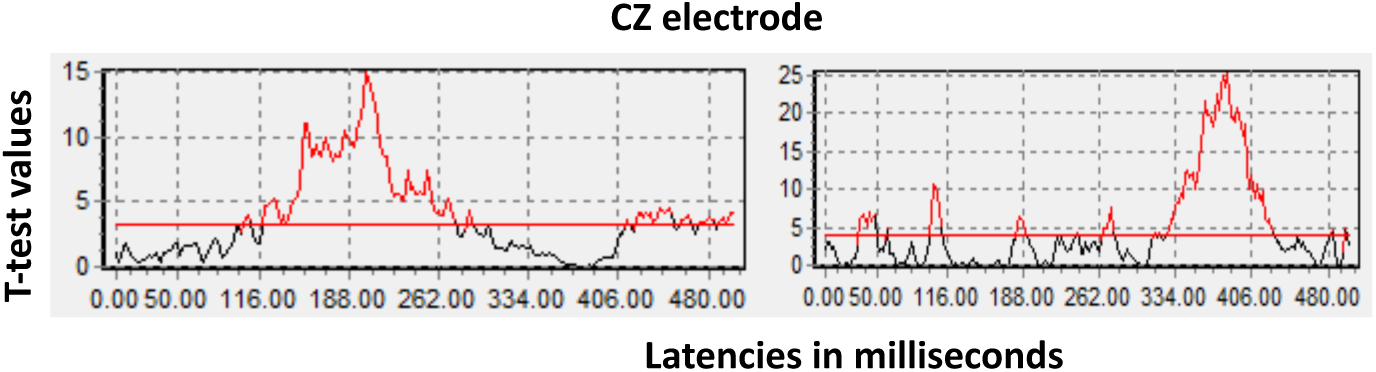
Left: t values for the tests of differences between Groups independently of Condition. Right: the t values tests for differences between Condition independently of Group. The read line indicates the statistical significant threshold (corrected for all electrodes and all times by a multivariate permutation test).

Note that the significant differences for Condition are in the range of the P300 or later, not in the scope of our study. For that reason, we focus all the further analysis on the incongruent condition, which is the condition which elicits inhibitory control. Nevertheless, henceforth we continue to report the full two-way analysis (Group × Condition), though concentrating on the Group Factor analyses.

### Analysis of the N1 component

We tested the N1 amplitudes with the repeated measures rMANOVA (Group X Condition), and examined the main effects and the interaction between them. The interaction and the factor Condition were not significant (p=0.23). However, the main effect of Group was significant with a Wilk’s Lambda=0.40, F(8,42)=2.97, p=0.009. A contrast between the two groups of patients was also significant with a Wilk’s Lambda=0.47, F(4,13)=1.2, p=0.003. Furthermore, with electrode-wise contrasts 13 electrode sites **F4, FC2, FC4, FC6, C2, C4, C6, CP2, O1, O2, Oz, PO3, PO4, PO7, PO8** retained significance. Note that the N1 at the O1 electrode followed the following pattern (See Figure 2): the amplitude of the PD rHuEPO group (−4.2 μV) was not different statistically from that of the HCs. On the other hand, the amplitude of the PD Control group (−1.2 μV) was significantly lower.

**Figure 2:**
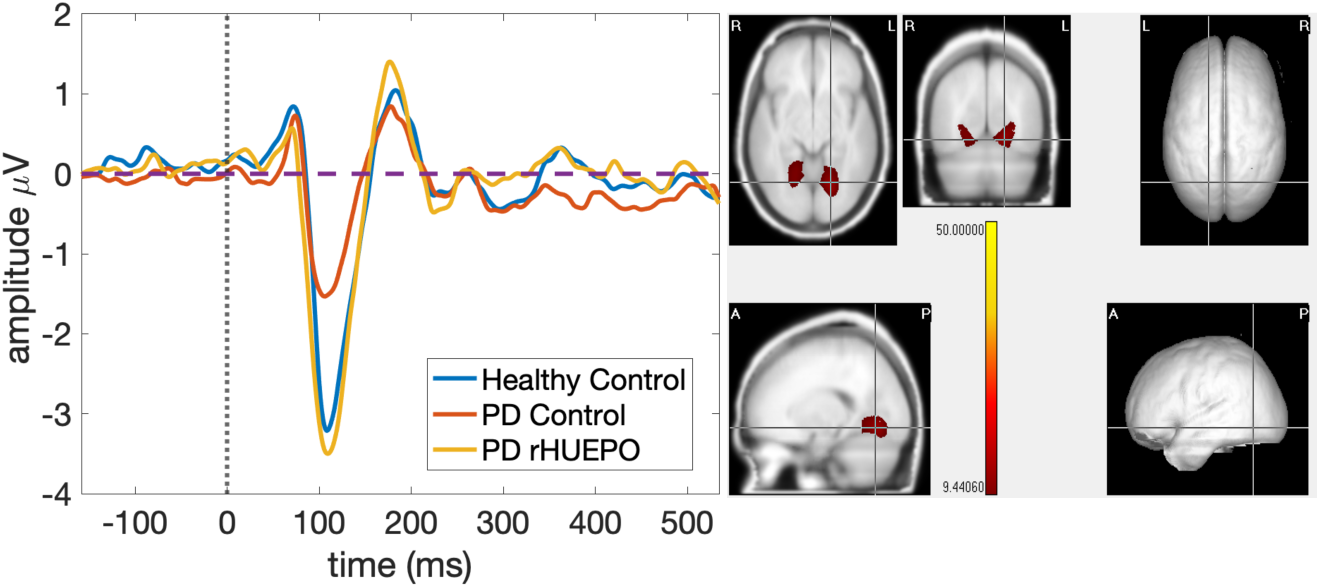
Left: the group average N1 waveform for each group in the window (80-180 msec.) in the electrode site O1 with the highest amplitude. The N1 peak was at 152 msec. Right: The Lingual Gyri are the sources of the N1 component according to AAL coordinates (X=92, Y=76, Z=172). The scale of statistical significance is self generated using the real values of the original data. All the voxels plotted were significant at p< 0.01).

The localization of the differences between the two Parkinson groups of this component are localized anatomically by means of the randomized nonparametric test for LORETA. This showed that the PD rHuEPO had a larger N1 component than the PD Control group at the p< 0.01 level (corrected for multiple comparisons) at the lingual gyri (See right side of Figure 2).

### Analysis of the N2 component

We tested the N2 amplitudes with the repeated measures rMANOVA (Group X Condition), and examined the interaction and the main effects. The interaction was not significant with a Wilk’s Lambda=0.43, F(6,44)=2.97 and the factor Condition was also not significant (p=0.323).

The main effect of Group (comparing **three groups**) was significant, F(2,24)=6.14, p=0.006, in seven fronto-central electrodes: Cz(F(2,24)=6.50, p=0.005),CPz (F(2,24)=4.43, p=0.02), CP1 (F(2,24)=5.9, p=0.008), CP2 (F(2,24)=5.6966, p=0.00945), C1(F(2,24)=3.6125, p=0.04251), C2 (F(2,24)=4.6242, p=0.02).

A contrast between the **two groups** of patients was also significant in fronto-central areas, the electrodes Cz (F(2,24)=4.43, p=0.002), CPz (F(2,24)=6.5, p=0.005) and FC1, FC2, C1, C2 (p<0.05). There were no significant differences between conditions or interaction between factors.

Note that the N2 grand average at the Cz electrode followed an opposite pattern than N1 (See Figure 4): the amplitude of the PD Control group (−2.10 μV) and healthy controls (−2.46 μV) was not different statistically. On the other hand, the amplitude of the PD rHuEPO group (−0.67 μV) was significantly lower than both of them. See Table 3 for details of amplitude and latencies of N2 in Cz.

**Figure 3:**
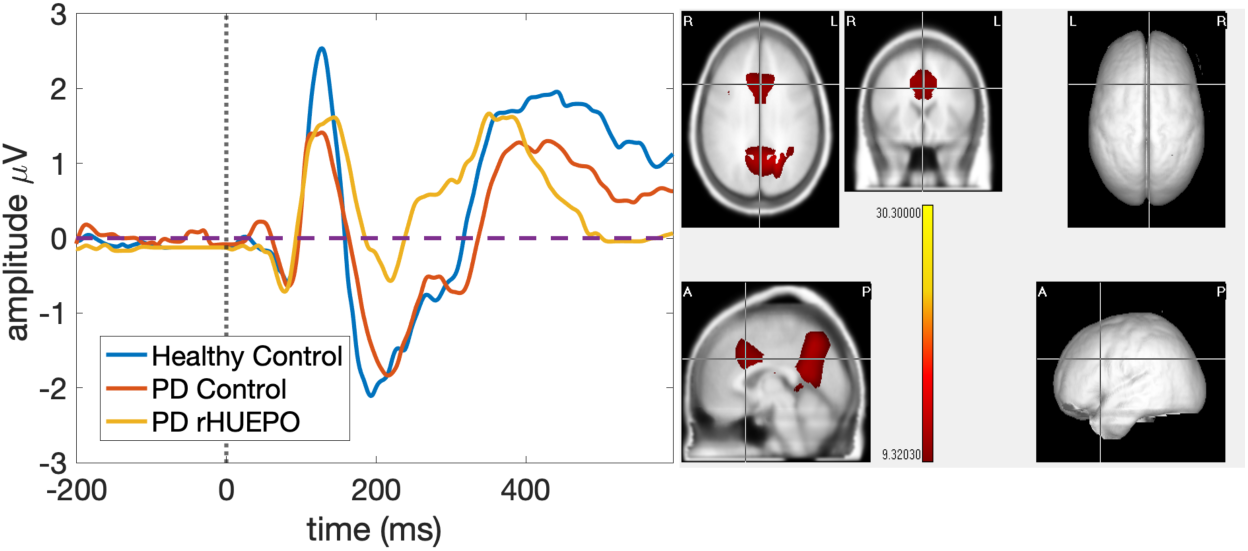
Left: the N2 waveform averaged by groups in the window (200-300 msec.) in the electrode site Cz with the highest amplitude. Note for the HC group the early 195 msec. latency and for both PD patients a later peak around 224 msec. Right: The N2 component showed maximal activation at middle cingulum and precuneus bilaterally (left located at X=92, Y=108, Z=156). To the right the localization of the precuneus left. The bicolour scale is showing all the signifcant values after Bonferroni correction and using permutations.

**Table 3:** The measures of amplitude and latency of the N2 component for the two conditions Congruent and Incongruent at the electrode Cz which exhibited the highest amplitude.

| Grupos | Amplitude ($\mu\text{V}$)<br>CONDITION | | Latency (msec.)<br>CONDITION | |
| --- | --- | --- | --- | --- |
|  | Cz-Cong | Cz-Incong | Cz-Cong | Cz-Incong |
| HCs | -2.4 | -2.45 | 195 | 199 |
| PD Control | -2.10 | -2.09 | 226 | 224 |
| PD rHuEPO | -0.56 | -0.67 | 224 | 223 |

**Figure 4:**
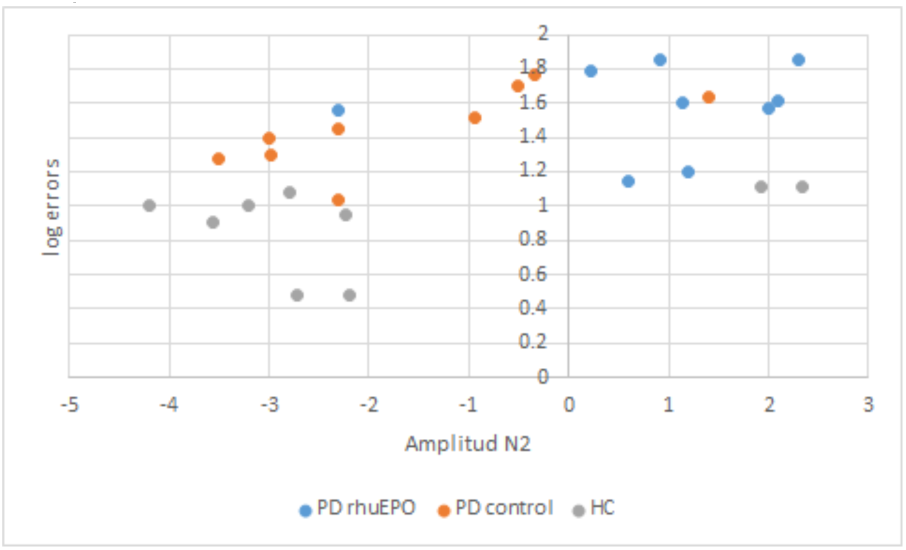
the plot of the N2 and log(errors) of the three groups. Note the variability of the data with 2 outliers of the HCs and 1 outlier of the PD Control group with positive amplitudes of N2.

The source analysis of the differences (**comparing the three groups**), for the N2 component, was localized anatomically by means of the LORETA randomized nonparametric test (p< 0.01 level corrected for multiple comparisons) at the the middle cingulum and precuneus bilaterally. See Figure 3, right side.

In order to know if the errors were related to the N2 amplitude, we select a linear mixed effect model, and carried out a repeated measures ANOVA log(errors) × Group × amplitudeN2. But the results were not significant for the interaction of log(errors) with the N2 amplitude, only the main effect for Group (p=0.001675). See Figure 4.

## Discussion

The current study was designed to examine if the novel rHuEPO neuroprotective compound, given to Parkinson patients in addition to their usual medication changed the amplitude of ERP components during an inhibitory control task.

The behavioural results were consisted with previous studies in PD patients in both the rHuEPO and PD Control groups. Both groups showed significantly increased reaction times and a higher number of errors to the incongruent stimuli during the performance of the flanker task as compared to age and education matched HCs. These higher error rates in PD controls are consistent with the proposal that the basal ganglia together with the anterior cingulate (Botvinick, Cohen, & Carter, 2004) participate in the monitoring of incongruence and error monitoring (Brázdil et al., 2002)(Michael Falkenstein, Christ, & Hohnsbein, 2000) which may be impaired in PD due to the dopamine deficiency (for a recent revision of how the progressive dopamine deficiency reduces striatal cholinergic interneuron activity see (McKinley et al., 2019).

It should be noted that we did not find the expected beneficial effect of rHuEPO on behavioural performance (RT and accuracy) in PD patients who received the neuroprotective agent as compared to those that only received the usual treatment. Rather, the differences between groups of patients were found in the ERP components. This is in accordance with our hypothesis that an overall behavioural response might be noisier than some of its time parsed substages. This suggests further studies to identify overt behavioural responses at similar short time scales as ERP components. On the other hand, as it sometimes happens with this type of clinical study the small sample size may lead to lack of power to detect subtle effects.

### Regarding the N1 component

This component reflects selective attention, linked to the basic characteristics of a stimulus, and also to the recognition of a specific visual pattern (Luck, Woodman, & Vogel, 2000). N1 amplitude also has been hypothesized to reflect sustained covert visual attention (Di Russo, Martinez, & Hillyard, 2003) being associated with the intensity of covert attention to the central target in the flanker task. In terms of spatial localization, the N1 amplitude is greater in occipital regions (Luck et al., 2000; Mangun & Hillyard, 1990). The neural sources of the N1 in Flanker tasks were located at the brain visual areas of the occipital cortex (Herrmann & Knight, 2001; Hillyard & Anllo-Vento, 1998; Luck et al., 2000). For example, Bokura (2001) using LORETA identified additional sources of the visual N1 in the occipito-temporal lobe (Bokura, Yamaguchi, & Kobayashi, 2001) and Zhang (2017) (Zhang, Brandt, Schrempf, Beste, & Stock, 2017) also localized N1 for Flanker source in extra-striate visual cortex. We thus expected the differences between PD groups to be localized on the scalp in the occipital electrodes and the sources to be in brain occipital areas.

This is what we found: the generators of N1, both in the scalp topography and using LORETA, in the visual areas of the occipital lobe of both hemispheres. The activation of the source for the PD patients who received rHuEPO was much larger that of the PD group who did not receive it. In fact, the response of the rHuEPO group became statistically indistinguishable from that of the HCs, suggestive of a possible neuroprotective effect of rHuEPO on the lingual gyrus, a region associated with the early and automatic processing of visual stimuli. In summary, our findings suggested an effect of rHuEPO on the visual attentional window in the early information-processing stage, thus enhancing the automatic processing of flankers regardless of their compatibility.

### Regarding the N2 component

The second component N2 has been found in several studies of inhibition using the Flanker task and its amplitude and latency was unaltered in medicated PD patients (for a review see (Seer et al., 2016)). Van Veen and Carter (Veen & Carter, 2002) used BESA source localization to study inhibition and response conflict in the Eriksen Flanker Task, determining that the N2 amplitude associated with incongruent trials can be explained by a dipole that is located in the ACC. Bokura et al. (Bokura et al., 2001) also conducted an experiment to understand the anatomical structures that are involved in N2 using a visual modality of the Flanker paradigm and LORETA which located the N2 generators at cingulate and the right lateral orbitofrontal cortex.

In our study, we found that the amplitude of N2 component for the PD control and HC groups, were statistically indistinguishable. But the N2 amplitude in the rHuEPO PD group was diminished with respect to the other two groups. These effects were topographically located, as expected, in the fronto-central areas, with neural generators of these differences localised to the posteromedial portion of the parietal lobe, the precuneus, a structure involved in the processing of perceptual ambiguities of stimuli (Cavanna & Trimble, 2006) and in the middle cingulate cortex, probably related to monitoring of conflict in the Flanker task (Enriquez-Geppert et al., 2013). In comparison with previous reports, we concur with Van Eimeren who found dysfunction of the default mode network and particularly deactivation of the posterior cingulate cortex and the precuneus (van Eimeren, Monchi, Ballanger, & Strafella, 2009) in PD relative to healthy controls, considering these changes in PD closely related to higher errors in executive tasks in PD compared with healthy controls.

However, in our study the striking decrease of the N2 produced by rHuEPO needs further research to find an adequate explanation.

### Behaviour versus ERPs

Contrary to our expectation, rHuEPO was not associated with a significant improvement in behavioural performance and did not influence the neural generators of the N2.

The ERP allows neural activity tracking on a millisecond time scale and represents a continuous measure of information processing, for that reason we selected the ERP to study a more refined measure of the process of inhibitory control.

This apparent contradiction between behavioural and electrophysiological results could be related to their different temporal course. Note that the inhibition is a complex process that can be automatically initiated in the first 100 msec. post-stimulus and extend its action through both automatic and controlled processes until 800 msec. Reaction time, on the other hand, started much later >400 milliseconds after the stimulus presentation, with a strong motor component to complete the response.

Therefore, the aim of our study is to use a flanker task to identify if rHuEPO improves automatic and controlled inhibitory control in PD patients and to locate the neural generators in these processes. This could be a first step in identifying an ERP biomarker for this type of cognitive process to be used in clinical trials.

### Limitations

Since this study was completed as part of a safety trial, the samples and the doses employed were small. This might also explain the lack of clear correlations with behaviour, for example reaction time with N2 amplitude. Thus, the results require confirmation with larger samples in future studies. However, the results highlighted the role of EEG source analysis and advantages of electrophysiology with its high temporal resolution and insensitivity to placebo effects, in identifying brain changes after an intervention such as rHuEPO.

## Conclusions

-We found that rHuEPO improved automatic inhibitory control in PD patients but did not improve behavioural performance.
-The differences between PD rHuEPO and PD Control groups was in the N1 component at the lingual gyrus. The differences between PD and healthy controls was on the N2 component in the cingulate and precuneus.
-Electrophysiology is potentially a useful tool for identifying effects of neuroprotective compounds on different stages of information processing.
-The components N1 and N2 as well as others like P3 should be further studied as possible biomarkers for the evaluation of neuroprotective drugs in Parkinson’s Disease.

## Data Availability

The tables with the behavioural performance (reaction time, hits and errors) and the N2 amplitude for the averaged time window (200-300 milliseconds) of the samples was submitted in the supplementary material 1. The raw and pre-processing EEG recordings in BrainVision format with all the individual and grand average potentials for group and condition can be available by request to

## Acknowledgments

The authors would like to thank to the Centro de Neurociencias de Cuba, specially to Valia Rodriguez and Indira Alvarez for the support during the recordings of the ERPs and the Centro Internacional de Restauracion Neurologica for the recruitment and neuropsychological evaluation of the patients. We are in debt with all the PD patients and their caretakers who volunteer to participated in our study.

## Supplementary Materials

The supplementary material 1 consisted in one excel table with the behavioural performance of the subjects during the Flanker task. Tab “Answers”: the hits, errors and non-answers in the congruent and incongruent condition. Tab “Reaction Time”: the mean and standard deviation (SD) of the hits, errors of each subject in each group for congruent and incongruent trials.

